# Effects of Cholesterol on Amyloid-induced Membrane Poration

**DOI:** 10.1101/2025.08.15.670631

**Authors:** Yanxing Yang, Bibhuti Shah, Maria Agapito, Andrew J. Nieuwkoop

## Abstract

The aggregation of amyloid peptides and their interactions with lipid membranes are central to the pathology of several neurodegenerative diseases. Using all-atom molecular dynamics simulations, we investigate how varying cholesterol concentrations (0%, 15%, and 30%) modulate amyloid-induced membrane poration. In cholesterol-free bilayers, pore formation was reproducibly observed in all simulations, whereas the presence of 15% cholesterol significantly reduced pore incidence, and 30% cholesterol entirely suppressed pore formation. Analysis revealed that cholesterol stiffens the bi-layer and strongly inhibits peptide-induced perturbations. Furthermore, cholesterol reduced the inter-leaflet mechanical coupling critical for transmembrane *β*-sheet formation, a key step in pore initiation. Our findings suggest that increasing cholesterol content protects membranes against amyloid-induced poration, providing a potential molecular basis for the observed reduction in amyloid toxicity in cholesterol-enriched environments. These results offer new insights into the complex role of membrane com-position in amyloid-related diseases and highlight cholesterol as a potential modulator of amyloid cytotoxicity.

## Introduction

Amyloid peptides are notorious for their central role in neurodegenerative diseases, including Alzheimer’s and Parkinson’s. ^1–3^ These peptides aggregate and interact with cellular membranes, leading to membrane disruption either via the formation of pores or through detergent-like mechanisms, ultimately resulting in cytotoxicity and cell death.^4–6^ Membrane poration, characterized by ion imbalance and increased permeability, has been extensively studied as a key pathogenic mechanism of amyloid-induced cytotoxicity.^7–9^

Computational simulations, particularly all-atom molecular dynamics (MD), have provided crucial atomic-level insights into these damage pathways, elucidating the structural and dynamic interactions between amyloid peptides and lipid bilayers. ^10,11^ Recent MD studies successfully captured spontaneous pore formation initiated by small amyloid-like peptide aggregates, highlighting significant lipid perturbations during membrane penetration. ^4,12^

Cholesterol is a major component of eukaryotic membranes, critically influencing their fluidity, thickness, and mechanical properties.^13–15^ Elevated cholesterol levels, particularly associated with aging, have been observed clinically in populations susceptible to Alzheimer’s disease.^16,17^ Interestingly, the impact of cholesterol on amyloid pathology is controversial and complex. ^18^ While some studies suggest cholesterol promotes amyloid aggregation and toxicity,^13,19^ others highlight cholesterol’s potential protective roles in modulating membrane structure and mitigating membrane damage.^15,20,21^

Specifically, cholesterol may significantly influence the propensity of amyloid peptides to insert into lipid membranes, affecting the overall stability of membrane structures upon peptide binding. ^22,23^ Recent computational studies have begun exploring these interactions. In particular, cholesterol-enriched membranes display increased order in lipid tails, leading to a reduced susceptibility to deformation and insertion by misfolded or aggregating proteins.^24^ In atomistic MD simulations, the presence of cholesterol has been shown to limit the hydrophobic mismatch and stabilize the membrane against the disruptive forces exerted by peptides. ^25^ Additionally, the presence of cholesterol can alter the local curvature stress and modulate the dynamics of lipid packing defects, which are often exploited by proteins to anchor themselves to membranes.^26^ These findings are consistent with experimental observations and support a growing consensus that cholesterol confers membrane resilience by inhibiting peptide aggregation at the membrane surface and reducing bilayer penetration. However, there remains a substantial gap in understanding precisely how varying cholesterol concentrations modulate membrane vulnerability to amyloid-induced damage at the molecular level.

In our previous work, we demonstrated via all-atom MD simulations the spontaneous aggregation of amphipathic peptides into *β*-sheets on lipid membranes and their subsequent insertion and pore formation, elucidating crucial atomic interactions governing these processes.^4^ Here, we significantly expand upon our earlier results by systematically varying the cholesterol concentration within lipid membranes (0%, 15%, and 30% cholesterol) and performing extensive MD simulations to probe its influence on amyloid-induced membrane poration.

Our results unequivocally demonstrate that the incorporation of cholesterol substantially inhibits pore formation. Specifically, spontaneous pore formation was consistently observed in cholesterol-free membranes, occurred rarely (one in five simulations) at intermediate cholesterol content (15%), and was entirely suppressed at higher cholesterol concentrations (30%). Furthermore, we observed that cholesterol markedly reduces the inter-leaflet interactions, effectively mitigating membrane perturbations induced by amyloid peptides. This is the first computational study providing direct atomic-scale evidence of cholesterol’s protective role by hindering amyloid insertion and subsequent pore formation, thereby suggesting a mechanistic explanation for the potential reduction of amyloid cytotoxicity in cholesterol-enriched membranes.

These findings offer significant insights into cholesterol’s dual role in amyloid diseases, potentially guiding therapeutic strategies aiming to modulate membrane cholesterol levels to mitigate amyloid-induced cellular damage. This work underscores the complexity and importance of membrane composition in amyloid pathophysiology, setting a foundation for future experimental and computational investigations.

## Methodology

In this study, we employed an amphipathic peptide with alternating non-polar (phenylalanine, F) and charged (lysine, K; glutamic acid, E) residues: Ac-(FKFE)_2_-NH_2_ (Figure 1a). This peptide is well-documented to self-assemble into cross-*β* structures, as demonstrated by both computational and experimental studies. ^27–29^ Our previous work further confirmed its ability to form *β*-sheets in membranous environments and to disrupt lipid bilayers within feasible simulation timescales.^4,30^

**Figure 1.**
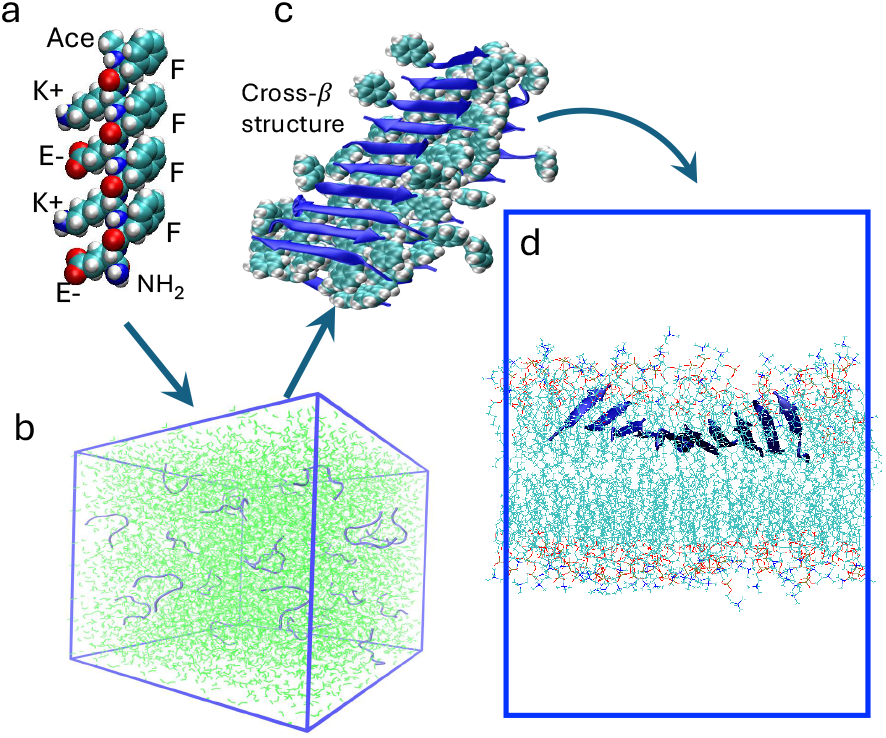
(a) Atomic structure of a single peptide, with side chains and termini labeled. (b) Simulation box containing 25 solvated peptides. (c) Cross-*β* structure formed by aggregated peptides, with non-polar side chains highlighted to emphasize hydrophobic interactions. (d) Schematic representation of the simulation setup showing the oligomer deposit interacting with a lipid bilayer.

### Preparation

Initially, 25 peptides were randomly positioned in a solvated cubic simulation box (Figure 1b). Following energy minimization and equilibration, the peptides self-assembled into a cross-*β* structure under the *NPT* ensemble (Figure 1c), as detailed in Ref.^29^ Subsequently, a *β*-sheet composed of 10 peptides was extracted and placed in a water box containing a lipid bilayer. To facilitate rapid insertion into the membrane core, the simulation box was constrained to 5.2 nm along the bilayer normal. After 100–300 ns of *NPT* simulation, the *β*-sheet bound irreversibly to the bilayer (Figure 1d).

### Simulation

The peptide-bound bilayer was placed in a 7*×*7*×*9 nm^3^ water box and solvated with 150 mM NaCl. Three bilayer compositions were modeled: 132 DMPC (CHL_0_, 0% cholesterol), 136 DMPC + 24 cholesterol (CHL_15_, 15%), and 112 DMPC + 48 cholesterol (CHL_30_, 30%). For each system, five 1-*µ*s simulations were conducted with varying insertion locations and orientations of the *β*-sheet to obtain statistical reliability. Additionally, three 100-ns control simulations without the peptide were performed for each bilayer composition.

### Protocol

Lipid bilayers were built using CHARMM-GUI^31–33^ with an initial box size of 7*×*7*×*12 nm^3^ and solvated using TIP3P water. Equilibration was performed in two stages using CHARMM-GUI protocols: three 5-ps *NV T* simulations with decreasing restraints, followed by three 5-ps *NPT* simulations with the same scheme.

All simulations were run using GROMACS-2022^34^ with the CHARMM36m force field. ^35^ The leapfrog integrator with a 2 fs time step was used. Temperature was maintained at 325 K via the Nosé-Hoover thermostat (*τ*_*T*_ = 1 ps),^36,37^ and pressure was maintained at 1 bar using the Parrinello-Rahman barostat (*τ*_*p*_ = 5 ps).^38^ Non-bonded interactions were calculated with a 1.2 nm cutoff and Verlet neighbor lists. Electrostatics were treated using the smooth particle mesh Ewald (PME) method with a grid spacing of 0.12 nm and a real-space cutoff of 1.2 nm.^39^

### Analysis

The secondary structure of peptides was analyzed using the STRIDE algorithm implemented in VMD.^40^ Intermolecular contacts between chemical groups were calculated using the GROMACS suite, with a cutoff distance of 0.4 nm to distinguish contact from non-contact states. Density distributions and diffusion coefficients were also evaluated using GROMACS tools. The number of water molecules (*N*_water_) in specific regions was estimated using in-house Python scripts based on the MDAnalysis library. ^41,42^ The lipid tail order parameter (*S*_*CC*_) was computed using custom Python code developed with the LiPyphilic toolkit.^43^

## Results and discussion

Poration of the lipid bilayer occurs when the bound *β*-sheet (induced phase) rotates and inserts into the membrane (poration phase), leading to increased bilayer permeability (Figure 2a,b). This transition is characterized by a sharp increase in the number of water molecules, *N*_water_, within the hydrophobic core of the bilayer (Figure 2c). Specifically, *N*_water_ remains below 50 in the absence of poration, but rises dramatically—up to approximately 400—when the *β*-sheet penetrates the membrane. Poration events were observed in all five 1-*µ*s replicates for the CHL_0_ system, indicating a high propensity for membrane disruption in the absence of cholesterol. In contrast, only one out of five replicates showed poration in the CHL_15_ system, and no poration events were observed in any of the five simulations for the CHL_30_ system, as summarized in Table 1. These results demonstrate a clear inverse correlation between cholesterol concentration and membrane poration by *β*-sheet aggregates. The presence of cholesterol appears to stabilize the bilayer structure and suppress peptide insertion, effectively inhibiting the formation of water-filled pores within the hydrophobic core. This trend is further supported by the suppressed water influx in the CHL_15_ and CHL_30_ systems, as shown in Figure 2c, which reflect reduced bilayer disruption in cholesterol-rich environments.

**Table 1.** Summary of poration events, corresponding maximum number of water molecules (*N*_water_) in the bilayer core for each system, and the lateral diffusion coefficient of the peptides moving in the bilayer.

| System | Poration Events (out of 5) | Maximum $N_{\text{water}}$ | Lateral Diffusion ($\text{nm}^2\mu\text{s}^{-1}$) |
| --- | --- | --- | --- |
| $\text{CHL}_0$ | 5 / 5 | $\sim 400$ | $1.83 \pm 0.27$ |
| $\text{CHL}_{15}$ | 1 / 5 | $\sim 350$ | $1.19 \pm 0.23$ |
| $\text{CHL}_{30}$ | 0 / 5 | $< 50$ | $0.99 \pm 0.24$ |

**Figure 2.**
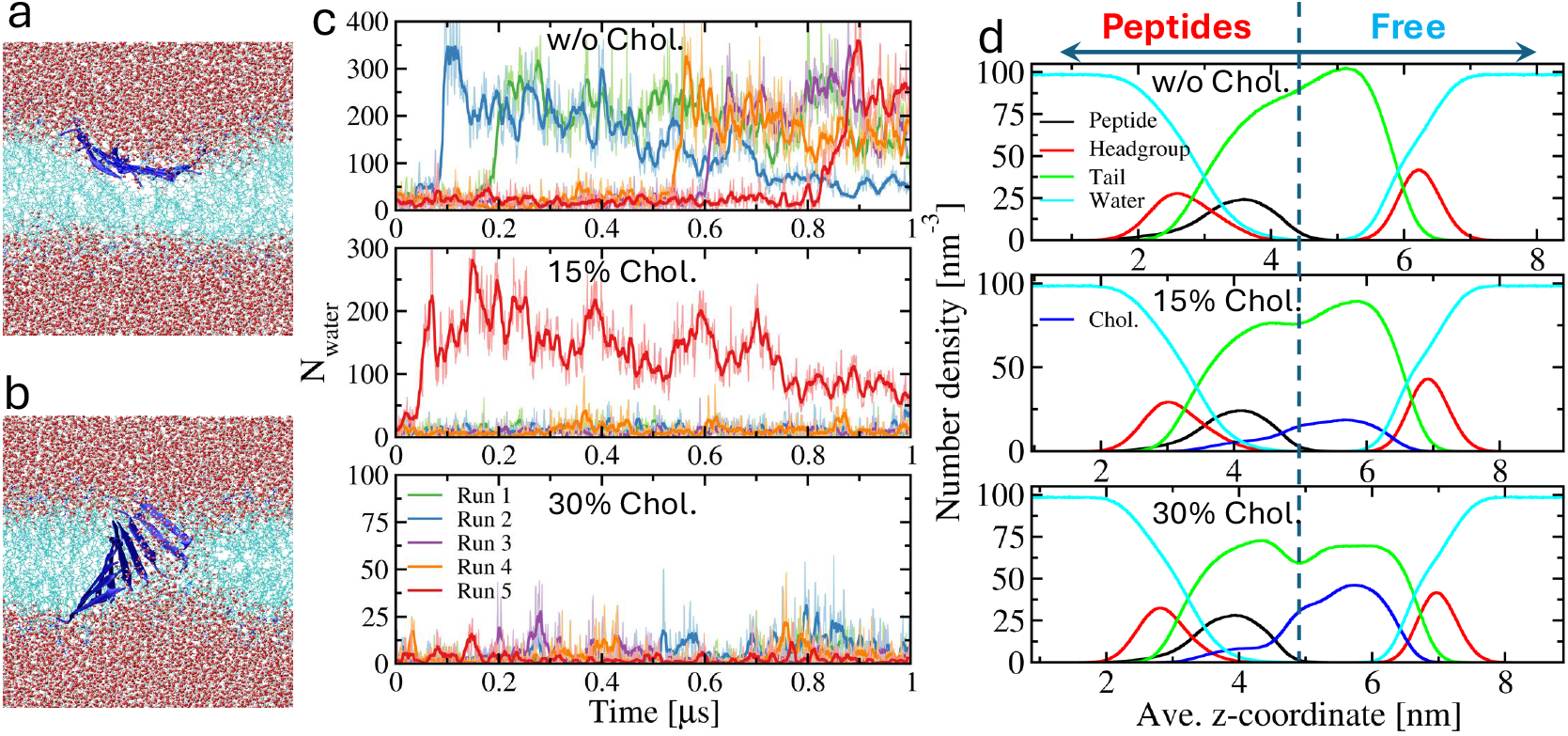
Cholesterol dependence of lipid bilayer poration. Visual representation of the *β* sheet (in blue) (a) bound to the bilayer surface and (b) penetrating the bilayer. (c) Time evolution of the number of water molecules in the hydrophobic core of the bilayer for CHL_0_, CHL_15_, and CHL_30_ systems (thin lines), with 10-point moving averages overlaid (thick lines). Atom number density profiles of key chemical groups along the bilayer normal (z-axis) during the induction phase. Data are centered at the bilayer midplane (vertical dashed line). Peptides bind to the leaflet on the left side of the midplane, while the opposite leaflet remains peptide-free.

The binding of peptides to a single leaflet of the bilayer induces significant asymmetry in membrane structure, as previously reported.^4^ This is clearly reflected in the number density profiles of lipid tail and cholesterol groups shown in Figure 2c. In the CHL_0_ and CHL_15_ systems (top two panels), the lipid tail densities (green lines) display a marked asymmetry between the peptide-bound and free leaflets, with a higher density observed in the latter. This redistribution suggests compression of the free leaflet, rendering it more susceptible to peptide-induced damage. In contrast, the CHL_30_ system (bottom panel) shows a more uniform tail distribution, with the free leaflet exhibiting a broad plateau rather than the distinct hump seen in the lower cholesterol systems. This plateau indicates a more stable and compact lipid organization, correlating with increased resistance to peptide-induced disruption. Furthermore, cholesterol (blue lines) also shows asymmetric localization, particularly in CHL_30_, where the majority resides in the free leaflet. This redistribution suggests that cholesterol preferentially migrates away from the perturbed, peptide-bound leaflet, contributing to the structural integrity of the opposing side. A previous study also supports this observation, reporting that peptide–membrane interactions can drive asymmetric cholesterol redistribution.^44^ Besides, The lateral diffusion coefficients of the peptide aggregates within the membrane exhibit a clear dependence on cholesterol concentration – see Table 1. In the CHL_0_ system, which lacks cholesterol, peptides diffuse more freely with an average diffusion coefficient of 1.83 nm^2^*µ*s^*−*1^. As cholesterol content increases, peptide mobility decreases significantly: 1.19 nm^2^*µ*s^*−*1^ in CHL_15_ and 0.99 nm^2^*µ*s^*−*1^ in CHL_30_. This reduction in diffusivity is consistent with the increased rigidity and packing density of cholesterol-rich bilayers, which hinder peptide movement and may further contribute to the suppression of poration events.^45^

To elucidate the role of hydrophobic interactions between lipid acyl chains and the nonpolar sidechains of the peptides, we quantified the average number of hydrophobic contacts between carbon atoms of the DMPC acyl chains and phenylalanine sidechains—see Figure 3a. Carbon atoms from cholesterol molecules residing at equivalent depths in the bilayer were also included in the analysis, and the carbon numbering scheme is illustrated in the inset. In cholesterol-free bilayers, the number of hydrophobic contacts increases progressively from the glycerol backbone (C1) toward the terminal methyl groups (C14), consistent with deeper insertion of the non-polar sidechains into the bilayer core.

**Figure 3.**
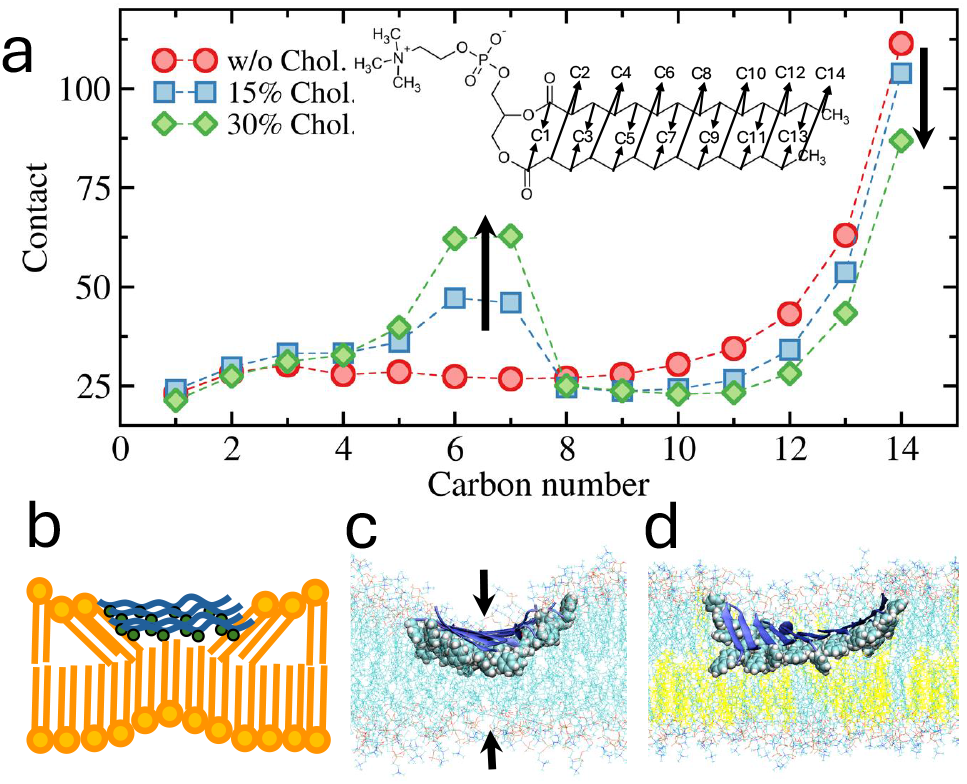
(a) Average number of hydrophobic contacts between the carbon atoms of DMPC acyl chains and the non-polar sidechains of the peptides. Carbon atoms from cholesterol molecules located at equivalent bilayer depths are also included in the analysis. The carbon numbering scheme is illustrated in the inset. (b) Schematic representation of the interactions between the non-polar sidechains of the peptides and lipid acyl chains. Representative configurations of the peptide–bilayer complexes in the (c) absence and (d) presence of cholesterol, which is highlighted in yellow.

In cholesterol-containing bilayers, while the overall depth-dependent trend is maintained, distinct changes emerge. Notably, the number of contacts involving the terminal tail carbons (C10–C14) decreases as cholesterol concentration increases. In contrast, a localized increase in contacts is observed at intermediate chain positions (C5–C7), where the interaction is notably enhanced in the presence of cholesterol, as evidenced by the contact “humps” in Figure 3a. This pattern suggests that cholesterol modulates the depth and spatial distribution of hydrophobic contacts within the membrane.

Mechanistically, these observations can be interpreted based on peptide orientation and membrane deformation. As depicted in Figure 3b, the non-polar sidechains are presented on the *β*-sheet surface facing the bilayer interior. Hydrophobic mismatch arises due to differences between the hydrophobic length of the peptide surface and the bilayer thickness. This mismatch is partially relieved by bilayer deformation, involving (1) tilting of lipid tails in the peptide-bound leaflet and (2) upward bending or “pulling up” of the free leaflet. The lipids from the bound leaflet primarily contact the edge residues of the *β*-sheet, whereas lipids from the free leaflet contribute to hydrophobic coverage over the central region.

This bilayer deformation, while compensating for mismatch, also reduces the local bilayer thickness, especially in the absence of cholesterol–see Figure 3c. In the presence of cholesterol, however, these effects are mitigated as shown in Figure 3d. Cholesterol preferentially accumulates in the free leaflet and increases local order and rigidity. As a result, the upward bending of the free leaflet is suppressed, leading to reduced contacts with the terminal acyl chain carbons (C10–C14) from that side. This stabilizing effect is consistent with the observed attenuation of bilayer thinning.

Interestingly, the reduction in deep-tail contacts is partially compensated by enhanced contacts with mid-chain carbons (C5–C7) in the bound leaflet. This suggests that cholesterol stabilizes the free leaflet while promoting adaptive interaction from the bound leaflet to maintain hydrophobic matching. Overall, cholesterol redistributes and fine-tunes the hydrophobic interactions across leaflets, stabilizing membrane structure and reducing local deformation induced by *β*-sheet-forming peptides.

To gain further insight into the structural perturbations induced by peptide insertion, we analyzed the lipid tail order parameter, *S*_*cc*_, which quantifies the orientational order of acyl chains and reflects membrane fluidity and packing defects.^46,47^ Two-dimensional maps of the 100 ns-averaged *S*_*cc*_, projected onto the membrane plane, are shown in Figure 4, with rows representing different leaflets and columns corresponding to varying cholesterol concentrations. Prior to analysis, the *β*-sheet was aligned at the center of the membrane, and no significant drift was observed during the 100 ns trajectory, allowing spatially resolved characterization of peptide-induced perturbations.

**Figure 4.**
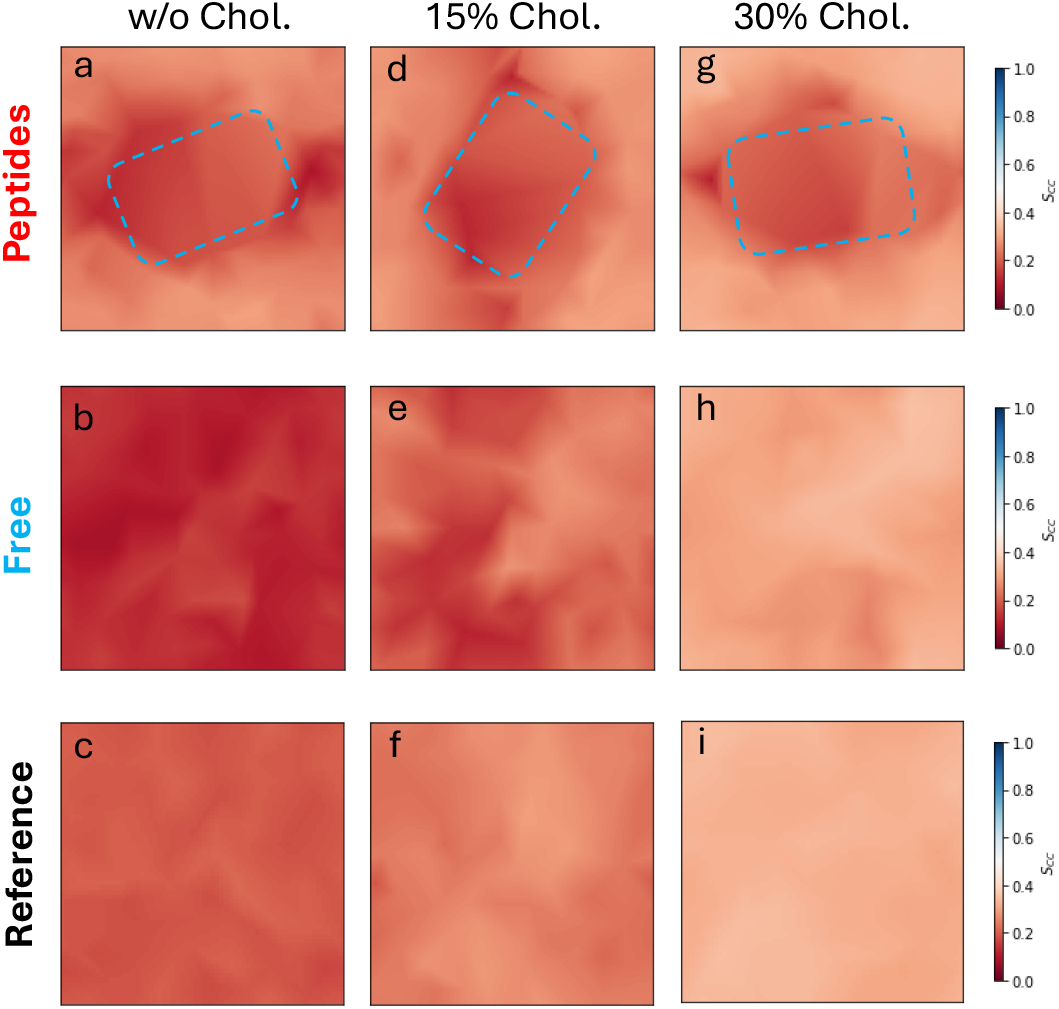
Two-dimensional maps of the 100 ns-averaged lipid tail order parameter (*S*_*cc*_) projected onto the membrane plane. The first row (a, d, g) corresponds to the peptide-bound leaflet, with the *β*-sheet position indicated by blue dashed lines. The second row (b, e, h) shows the free leaflet, and the third row (c, f, i) depicts the reference bilayers. Columns represent bilayers with (a–c) no cholesterol, (d–f) 15% cholesterol, and (g–i) 30% cholesterol.

In the absence of peptides (third row: panels c, f, i), the *S*_*cc*_ values are distributed homogeneously across the membrane and increase systematically with rising cholesterol content, consistent with the known ordering effect of cholesterol on lipid acyl chains. This provides a baseline for comparison against peptide-bound systems.

In contrast, the peptide-bound leaflets (first row: panels a, d, g) exhibit clear disruptions in lipid order near the *β*-sheet region, indicated by blue dashed lines. The *S*_*cc*_ is significantly reduced in the vicinity of the peptides, reflecting local disorder induced by peptide insertion. Importantly, this disruption is spatially confined and decays rapidly with increasing distance from the *β*-sheet center. This localized effect is consistent with our previous findings. ^4^

The free leaflet (second row: panels b, e, h) exhibits more extensive perturbation compared to the peptide-bound leaflet, particularly in the CHL_0_ system (Figure 4b), where a substantial reduction in *S*_*cc*_ extends over a broader region. This suggests that peptide insertion into one leaflet can indirectly disrupt the opposite leaflet, likely due to transbilayer stress coupling and hydrophobic mismatch. As cholesterol concentration increases, this effect is progressively attenuated. In the CHL_30_ system, the *S*_*cc*_ distribution in the free leaflet (Figure 4h) closely resembles that of the reference bilayer (Figure 4i), indicating a return to a more ordered and resilient membrane state.

These observations support the interpretation illustrated in Figure 3b: peptides induce local thinning and deformation in the peptide-bound leaflet, which in turn causes upward displacement and disordering of the free leaflet. Cholesterol, preferentially enriched in the free leaflet, rigidizes the membrane and mitigates this deformation, thereby confining the perturbation to the peptide-bound region. Moreover, cholesterol likely improves hydrophobic matching in the bound leaflet, reducing the need for compensatory deformation in the opposite leaflet.

In summary, *S*_*cc*_ mapping reveals that peptide-induced membrane disorder is localized and cholesterol-dependent. While peptide insertion strongly disrupts the immediate vicinity in the bound leaflet, cholesterol dampens the long-range propagation of this effect across the bilayer, reinforcing membrane resilience.

## Conclusion

In this study, we employed all-atom molecular dynamics simulations to investigate how cholesterol modulates amyloid-induced membrane perturbation and pore formation in lipid bilayers. Our simulations reveal a clear cholesterol-dependent effect that robust pore formation consistently occurred in cholesterol-free bilayers was significantly reduced at 15% cholesterol, and was completely suppressed at 30% cholesterol. These results underscore cholesterol’s critical role in stabilizing membrane structure against peptide-induced disruption.

Beyond inhibiting pore formation, cholesterol altered the inter-leaflet coupling of the bilayer. In cholesterol-free and low-cholesterol membranes, binding of amyloid-like peptides to one leaflet strongly distorted the opposing leaflet, leading to asymmetric density profiles and a mechanically fragile bilayer. By contrast, in cholesterol-rich membranes, inter-leaflet perturbation was substantially reduced, and lipid packing in the free leaflet became more compact and resilient. Analysis of lipid order parameters further confirmed that cholesterol enhances local ordering of lipid tails and reduces peptide-induced disorder.

Together, these findings provide molecular-level evidence that cholesterol protects mem-branes from amyloid-induced damage by reinforcing lipid packing, limiting inter-leaflet communication of stress, and thereby preventing pore formation. Besides the mechanistic insight into the protective effect of cholesterol on membrane integrity, this work also highlights the importance of bilayer composition in modulating amyloid cytotoxicity. Such insights could inform therapeutic strategies that target membrane composition to mitigate amyloid-related pathologies.

